# Contributions of plasmid p1AB5075-encoded antibiotic resistance genes to multidrug resistance of *Acinetobacter baumannii* AB5075

**DOI:** 10.64898/2026.03.29.715119

**Authors:** Orlaith Plunkett, Anna S. Ershova, Kristina Schauer, Carsten Kröger

## Abstract

Infections with multidrug resistant *Acinetobacter baumannii* are considered a threat to human and animal health. The widely studied *A. baumannii* strain AB5075 displays a high degree of antibiotic resistance. In this study, we experimentally validated that antibiotic resistance is largely mediated by resistance genes located on plasmid p1AB5075. We used a p1AB5075-deficient AB5075 strain to assess individual contributions of p1AB5075-encoded antibiotic resistance genes by ectopically (over-)expressing each gene in the Δp1AB5075 background. By determining individual contributions of seven p1AB5075-encoded antibiotic resistance genes, we show individual and overlapping roles of genes for aminoglycoside resistance and uncover the importance of extended-spectrum ß-lactamase *bla*_*GES-11*_ for cephalosporin resistance in *A. baumannii* AB5075. We discovered that aminoglycoside N-acetyltransferase *aaC(6’)-Ib3* (*aacA4*), which was considered a potential pseudogene in *A. baumannii* AB5075, to be functional providing broad resistance to gentamicin, kanamycin, amikacin, streptomycin and tobramycin when overexpressed in *A. baumannii* AB5075. Because p1AB5075 is transferrable to a wide range of environmental and clinical *A. baumannii* strains and non-*baumannii Acinetobacter* species, the relevance of our findings extends beyond *A. baumannii* AB5075.

## INTRODUCTION

Curing infections with multidrug resistant (MDR) microorganisms of humans and animals is one of the greatest medical challenges of the 21^st^ century. Among the most problematic bacteria that require investment into research and development of new antibiotics or alternative treatment strategies are carbapenem-resistant *Acinetobacter baumannii* (Tacconelli *et al*., 2018; Whiteway *et al*., 2022a; Sati *et al*., 2025). Infections with *A. baumannii* are increasingly difficult to clear because of a high degree of antimicrobial resistance, strain heterogeneity and reports of pan-drug resistant strains (Valcek *et al*., 2022a; Cain and Hamidian, 2023; Valcek *et al*., 2025). Understanding the evolution of antimicrobial resistance and resistance mechanisms may inform the development of new drugs and better treatment regimes. Early isolates that have been extensively used to study *A. baumannii* biology were typically sensitive to antibiotics including the widely studied strains *A. baumannii* ATCC17978 and ATCC19606 (Smith *et al*., 2007). Contemporary *A. baumannii* strains, including the widely studied *A. baumannii* AB5075, are likely more suitable representatives of current infections, and are often resistant to a large array of antibiotics (Jacobs *et al*., 2014). In *A. baumannii* AB5075, many antibiotic resistance genes (ARGs) are located on plasmid p1AB5075 within “Resistance Island 2” (RI-2), but some also are found on the chromosome (e.g., *oxa*-23) (Jacobs *et al*., 2014; Gallagher *et al*., 2015). The plasmid p1AB5075 is the largest of three plasmids in *A. baumannii* AB5075 (83,160 bp), which was shown to be transferrable to other *A. baumannii* strains of environmental and clinical origin and to diverse non-*baumannii Acinetobacter* species (Jacobs *et al*., 2014; Gallagher *et al*., 2015; Nasser *et al*., 2024; Martz *et al*., 2025). The RI-2 locus is chiefly responsible for aminoglycoside (hetero-)resistance in *A. baumannii* AB5075, which is caused in part by a RecA-dependent resistance island amplification to up to 20-24 copies (Anderson *et al*., 2018). The RI-2 small RNA SrvS located upstream of *aadB* was also shown to be involved in virulent opaque (VIR-O) to avirulent translucent colony (AV-T) type switching, showing that p1AB5075-encoded genes are engaging in regulatory cross-talk between p1AB5075 and the chromosome (Anderson *et al*., 2020). In addition, another opaque subpopulation of AB5075 termed LSO (low switching opaque) is distinct from the VIR-O variant by reduced switching frequency compared to VIR-O. The number of RI2 copies (the LSO variant has only one copy, while the VIR-O variant has ≥2 copies) influences switching frequency between opaque and translucent variants through the copy number of the *srvS* gene (Anderson *et al*., 2020). RI-2 is mosaic and contains among other ARGs an extended-spectrum β-lactamase *bla*_GES-11_, which was first described in an *A. baumannii* strain isolated in France (Moubareck *et al*., 2009). Despite their biological importance, spontaneous loss of large plasmids, including p1AB5075, has been documented for *A. baumannii* (Valcek *et al*., 2025). Plasmid pAB3 has been reported to have been lost in *A. baumannii* ATCC17978, which caused up-regulation of the Type-6 Secretion System (Weber *et al*., 2015; Kröger *et al*., 2018), and loss of p1AB5075 in *A. baumannii* AB5075 resulted in sensitivity to amikacin and tobramycin and reduced resistance to chloramphenicol (Anderson *et al*., 2020; de Dios *et al*., 2022). Here, we report loss of p1AB5075 in *A. baumannii* AB5075 independent from the published reports (Anderson *et al*., 2020; de Dios *et al*., 2022). We used the Δp1AB5075 genetic background to characterise the antibiotic resistance profile for the p1AB5075-deficient strain and genetically dissect the contributions of seven individual p1AB5075-encoded antibiotic resistance genes for high level multidrug resistance.

## METHODS

### Bacterial strains and general growth conditions

*Acinetobacter baumannii* AB5075 and *Escherichia coli* TOP10 (Invitrogen) were maintained on lysogeny broth (Lennox, L-) agar plates (10 g/l tryptone, 5 g/l yeast extract, 5 g/l NaCl, 15 g/l agar) and grown over night in liquid L-broth, 10 g/l tryptone, 5 g/l yeast extract, 5 g/l NaCl) (Jacobs *et al*., 2014). Media contained tetracycline (12 μg/ml) when necessary to maintain plasmid pWH1266 (Hunger *et al*., 1990).

### Strain and plasmid constructions

ARGs were amplified from purified genomic DNA of *A. baumannii* AB5075 using DNA oligonucleotides listed in **Supplementary Table 1** with ∼20 bp overhangs complementary to the pWH1266 backbone to enable sequence and ligation independent cloning extract (SLiCE)-mediated cloning (Zhang *et al*., 2012). Polymerase chain reaction (PCR) amplification of DNA was performed using Verify™ polymerase (PCR Biosystems). Plasmid isolations and PCR purifications were carried out according to the manufacturer’s protocol using EasyPure Plasmid MiniPrep and PCR Purification Kits (TransGen Biotech). SLiCE cloning procedure was used to insert ARGs into the PCR-amplified pWH1266 backbone (Zhang *et al*., 2012). All plasmid construction steps were carried out in *E. coli* TOP10 cells. To ensure efficient and equal translation of ARG mRNAs, the forward primer contained an artificial AGGAGG ribosome binding site (RBS). The cloned genes were constitutively expressed from the β-lactamase (*bla*) promoter of pWH1266 (Hunger *et al*., 1990). Plasmids were re-isolated from *E. coli*, their insert sequence verified by Sanger sequencing (Eurofins), and AB5075 was subsequently transformed with the plasmids using natural transformation or electroporation (Godeux *et al*., 2020). Whole plasmid sequencing of pWH1266 was conducted by Azenta/GENEWIZ (GENEWIZ Germany GmbH) using Oxford Nanopore Technology (ONT) in a GridION with a FLO-MIN114 (R10.4.1) flow cell (GenBank accession no.: PV577797.1).

### Disk diffusion assays

All *A. baumannii* AB5075 strains were cultured overnight in cation-adjusted Mueller-Hinton broth (MH2, Merck/Millipore). Overnight cultures were adjusted to 0.5 McFarland standard, and each culture sample was swabbed thoroughly onto Mueller-Hinton agar plates using a sterile cotton swab. Plates were air-dried for 10 min before addition of antimicrobial susceptibility disks (Oxoid). Antibiotic disks contained amikacin (AK, 30 μg), tobramycin (TOB, 10 μg), kanamycin (K, 30 μg), gentamicin (CN, 10 μg), trimethoprim-sulfamethoxazole (SXT, 25 μg), streptomycin (S, 25 μg), meropenem (MEM, 10 μg), doripenem (DOR, 10 μg), imipenem (IMP 10 μg), vancomycin (VA, 30 μg), erythromycin (E, 15 μg), nitrofurantoin (F, 100 μg), tetracycline (TE, 30 μg), tigecycline (TGC 15 μg), ciprofloxacin (CIP, 5 μg), cefepime (FEP, 30 μg), cefoperazone (CFP, 30 μg), ceftazidime (CAZ, 30 μg), chloramphenicol (C, 30 μg), aztreonam (ATM, 30 μg), oxacillin (OX, 1 μg), ticarcillin (TIC, 75 μg), cefoxitin (FOX, 30 μg) and piperacillin/tazobactam (TZP, 110 μg). Plates were incubated over night at 37°C before measuring the diameters of inhibition zones.

### Broth microdilution assay

Broth microdilution assays were performed as per the Clinical & Laboratory Standards Institute (CLSI) guidelines for *Acinetobacter* to determine the Minimum Inhibitory Concentrations (MIC) for antibiotics. The assays were performed in 96-well plates. Ten µl of culture (0.5 McFarland) was added to each well except for the negative control (MH2 medium without bacteria). Plates were incubated at 37°C statically for 22 h before the plates were analysed visually.

## RESULTS

### Loss of p1AB5075 results in extensive sensitivity to multiple antibiotics

*Acinetobacter baumannii* AB5075 is considered a multidrug resistant strain with increased virulence and a contemporary representative of *A. baumannii* infections (Jacobs *et al*., 2014). Because *A. baumannii*, and especially *A. baumannii* AB5075, shows a high degree of phenotypic and genotypic heterogeneity (Chin *et al*., 2018; Whiteway *et al*., 2022b; Pérez-Varela *et al*., 2022; Valcek *et al*., 2022b; Cooper *et al*., 2024; Singh *et al*., 2025; Valcek *et al*., 2025), we whole-genome sequenced a number of *A. baumannii* AB5075 colonies after attempting to delete genes AB5075 by the suicide-vector method of Pokhrel *et al*. (Pokhrel *et al*., 2023). Serendipitously, this revealed that one of the sequenced colonies, where the wild-type genotype at the position of the attempted deletion was restored, had lost plasmid p1AB5075 but was otherwise wild-type—an event which had been independently reported before (de Dios *et al*., 2022). As p1AB5075 carries multiple antibiotic resistance genes as part of RI-2 (Gallagher *et al*., 2015), we set out to characterise the antibiotic resistance profile of the Δp1AB5075 strain and to investigate the individual contributions of ARGs on p1AB5075, which so far, with the exception of *cmlA* contributing to chloramphenicol resistance had not been experimentally verified gene-by-gene in *A. baumannii* AB5075 (de Dios *et al*., 2022). The predicted ARGs of p1AB5075 are shown in the context of p1AB5075 in **Figure 1** and their annotation is presented in **Table 1**. All ARGs except one, a predicted APH(3’)-VI family aminoglycoside O-phosphotransferase (*adh*/*ABUW_RS19420*), are located within RI-2 (**Figure 1A & B**). Antibiotic disk diffusion assays were performed to assess antibiotic resistance profiles of wild-type AB5075 and the Δp1AB5075 mutant strain to a suite of antibiotics. The Δp1AB5075 mutant strain showed widespread increased susceptibility including to all tested aminoglycosides (amikacin, gentamicin, kanamycin, streptomycin, tobramycin), but also to trimethoprim/sulfamethoxazole, aztreonam (monobactam) and ceftazidime (cephalosporin) **(Figure 2, Table 2)** highlighting the crucial role for p1AB5075 towards multi-drug resistance. No difference was observed to ciprofloxacin (fluoroquinolone), cefoperazone, cefepime (cephalosporins), doripenem, imipenem, meropenem (carbapenems), nitrofurantoin (nitrofuran), erythromycin (macrolide), vancomycin (glycopeptide), tetracycline, tigecycline (tetracyclines), cefoxitin (cephamycin), oxacillin, ticarcillin (β-lactams), chloramphenicol and the broad-spectrum β-lactam antibiotic/inhibitor combination piperacillin/tazobactam. We noted a difference in cephalosporin resistance, where wild-type and Δp1AB5075 strains were resistant to cefoperazone and cefepime but not ceftazidime indicating that ceftazidime resistance is mediated by Δp1AB5075 (**Figure 2**). Resistance to carbapenems did not change which is mediated by a chromosomally encoded *oxa*-23 gene (Intorcia *et al*., 2024). Resistance to chloramphenicol was not altered either, likely to the presence of additional chloramphenicol resistance genes *craA* and *cpxE* (*ABUW_0982*) (Roca *et al*., 2009; Karalewitz and Miller, 2018).

**Table 1:**
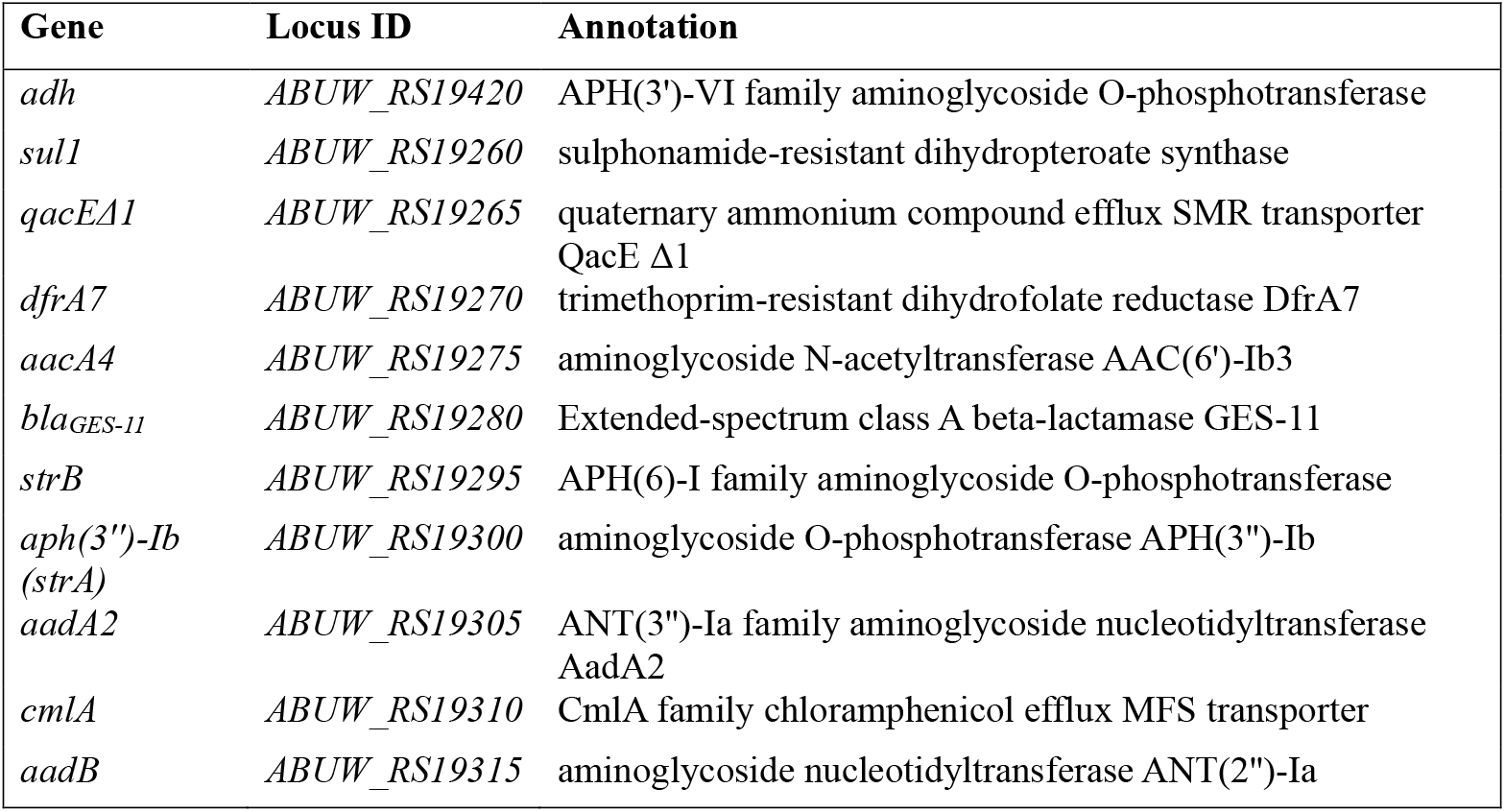
Annotation of antibiotic resistance genes of p1AB5075.

**Table 2:**
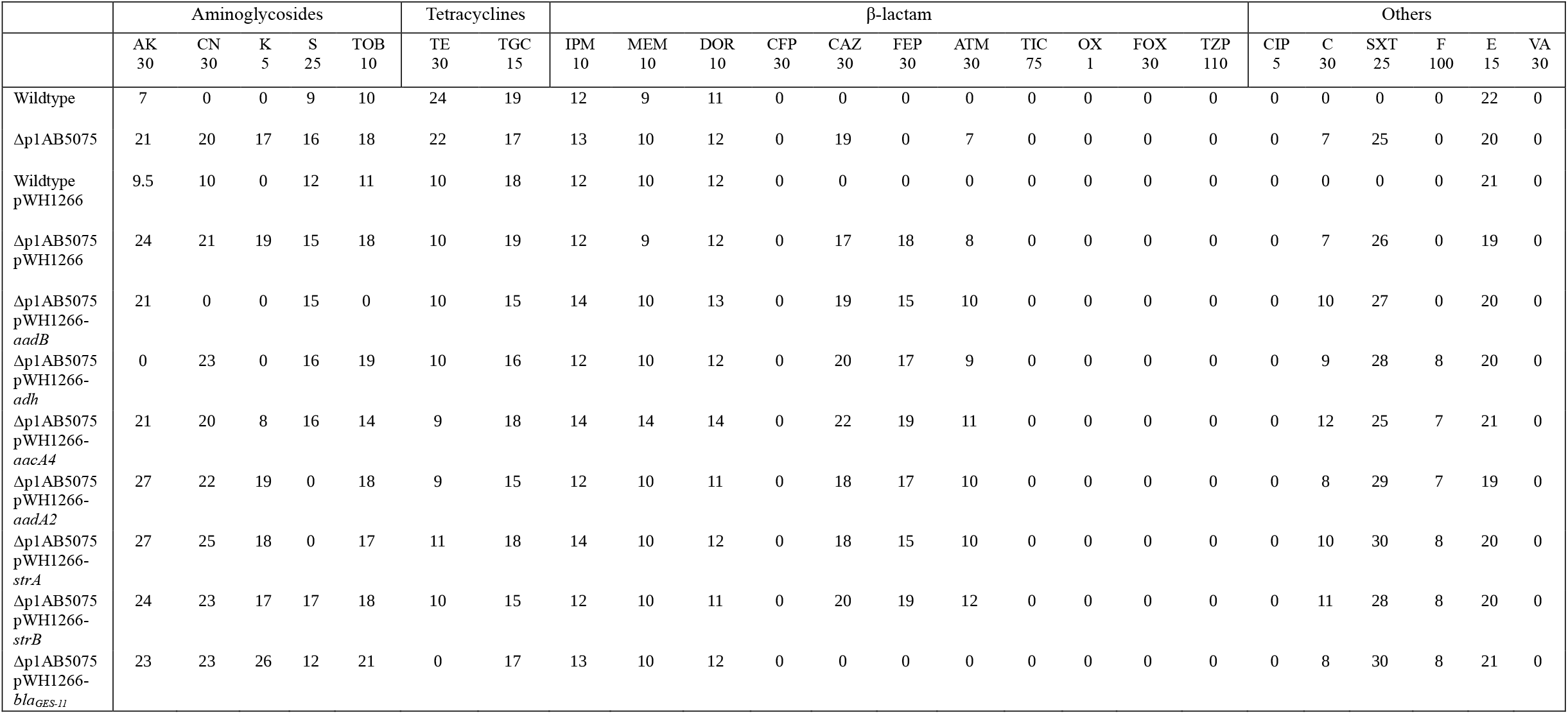
Average of inhibition zones of AB5075 mutants measured in mm (n = 3) determined by Kirby-Bauer disk diffusion assays. AK=amikacin, CN=gentamicin, K=kanamycin, S=streptomycin, TOB=tobramycin, TE=tetracycline, TGC=tigecycline, IPM=imipenem, MEM=meropenem, DOR=doripenem, CFP=, CAZ=ceftazidime, FEP=cefepime, ATM=aztreonam, TIC=ticarcillin, OX=oxacillin, FOX=cefoxitin, TZP=piperacillin/tazobactam, CIP=ciprofloxacin, C=chloramphenicol, SXT=trimethoprim/sulfamethoxazole, F=nitrofurantoin, E=erythromycin, VA=vancomycin.

|  | Aminoglycosides |  |  |  |  | Tetracyclines |  | β-lactam |  |  |  |  |  |  |  |  |  |  | Others |  |  |  |  |  |
| --- | --- | --- | --- | --- | --- | --- | --- | --- | --- | --- | --- | --- | --- | --- | --- | --- | --- | --- | --- | --- | --- | --- | --- | --- |
|  | AK<br>30 | CN<br>30 | K<br>5 | S<br>25 | TOB<br>10 | TE<br>30 | TGC<br>15 | IPM<br>10 | MEM<br>10 | DOR<br>10 | CFP<br>30 | CAZ<br>30 | FEP<br>30 | ATM<br>30 | TIC<br>75 | OX<br>1 | FOX<br>30 | TZP<br>110 | CIP<br>5 | C<br>30 | SXT<br>25 | F<br>100 | E<br>15 | VA<br>30 |
| Wildtype | 7 | 0 | 0 | 9 | 10 | 24 | 19 | 12 | 9 | 11 | 0 | 0 | 0 | 0 | 0 | 0 | 0 | 0 | 0 | 0 | 0 | 0 | 22 | 0 |
| Δp1AB5075 | 21 | 20 | 17 | 16 | 18 | 22 | 17 | 13 | 10 | 12 | 0 | 19 | 0 | 7 | 0 | 0 | 0 | 0 | 0 | 7 | 25 | 0 | 20 | 0 |
| Wildtype<br>pWH1266 | 9.5 | 10 | 0 | 12 | 11 | 10 | 18 | 12 | 10 | 12 | 0 | 0 | 0 | 0 | 0 | 0 | 0 | 0 | 0 | 0 | 0 | 0 | 21 | 0 |
| Δp1AB5075<br>pWH1266 | 24 | 21 | 19 | 15 | 18 | 10 | 19 | 12 | 9 | 12 | 0 | 17 | 18 | 8 | 0 | 0 | 0 | 0 | 0 | 7 | 26 | 0 | 19 | 0 |
| Δp1AB5075<br>pWH1266-<br>*aadB* | 21 | 0 | 0 | 15 | 0 | 10 | 15 | 14 | 10 | 13 | 0 | 19 | 15 | 10 | 0 | 0 | 0 | 0 | 0 | 10 | 27 | 0 | 20 | 0 |
| Δp1AB5075<br>pWH1266-<br>*adh* | 0 | 23 | 0 | 16 | 19 | 10 | 16 | 12 | 10 | 12 | 0 | 20 | 17 | 9 | 0 | 0 | 0 | 0 | 0 | 9 | 28 | 8 | 20 | 0 |
| Δp1AB5075<br>pWH1266-<br>*aacA4* | 21 | 20 | 8 | 16 | 14 | 9 | 18 | 14 | 14 | 14 | 0 | 22 | 19 | 11 | 0 | 0 | 0 | 0 | 0 | 12 | 25 | 7 | 21 | 0 |
| Δp1AB5075<br>pWH1266-<br>*aadA2* | 27 | 22 | 19 | 0 | 18 | 9 | 15 | 12 | 10 | 11 | 0 | 18 | 17 | 10 | 0 | 0 | 0 | 0 | 0 | 8 | 29 | 7 | 19 | 0 |
| Δp1AB5075<br>pWH1266-<br>*strA* | 27 | 25 | 18 | 0 | 17 | 11 | 18 | 14 | 10 | 12 | 0 | 18 | 15 | 10 | 0 | 0 | 0 | 0 | 0 | 10 | 30 | 8 | 20 | 0 |
| Δp1AB5075<br>pWH1266-<br>*strB* | 24 | 23 | 17 | 17 | 18 | 10 | 15 | 12 | 10 | 11 | 0 | 20 | 19 | 12 | 0 | 0 | 0 | 0 | 0 | 11 | 28 | 8 | 20 | 0 |
| Δp1AB5075<br>pWH1266-<br>*bla<sub>GES-11</sub>* | 23 | 23 | 26 | 12 | 21 | 0 | 17 | 13 | 10 | 12 | 0 | 0 | 0 | 0 | 0 | 0 | 0 | 0 | 0 | 8 | 30 | 8 | 21 | 0 |

**Figure 1:**
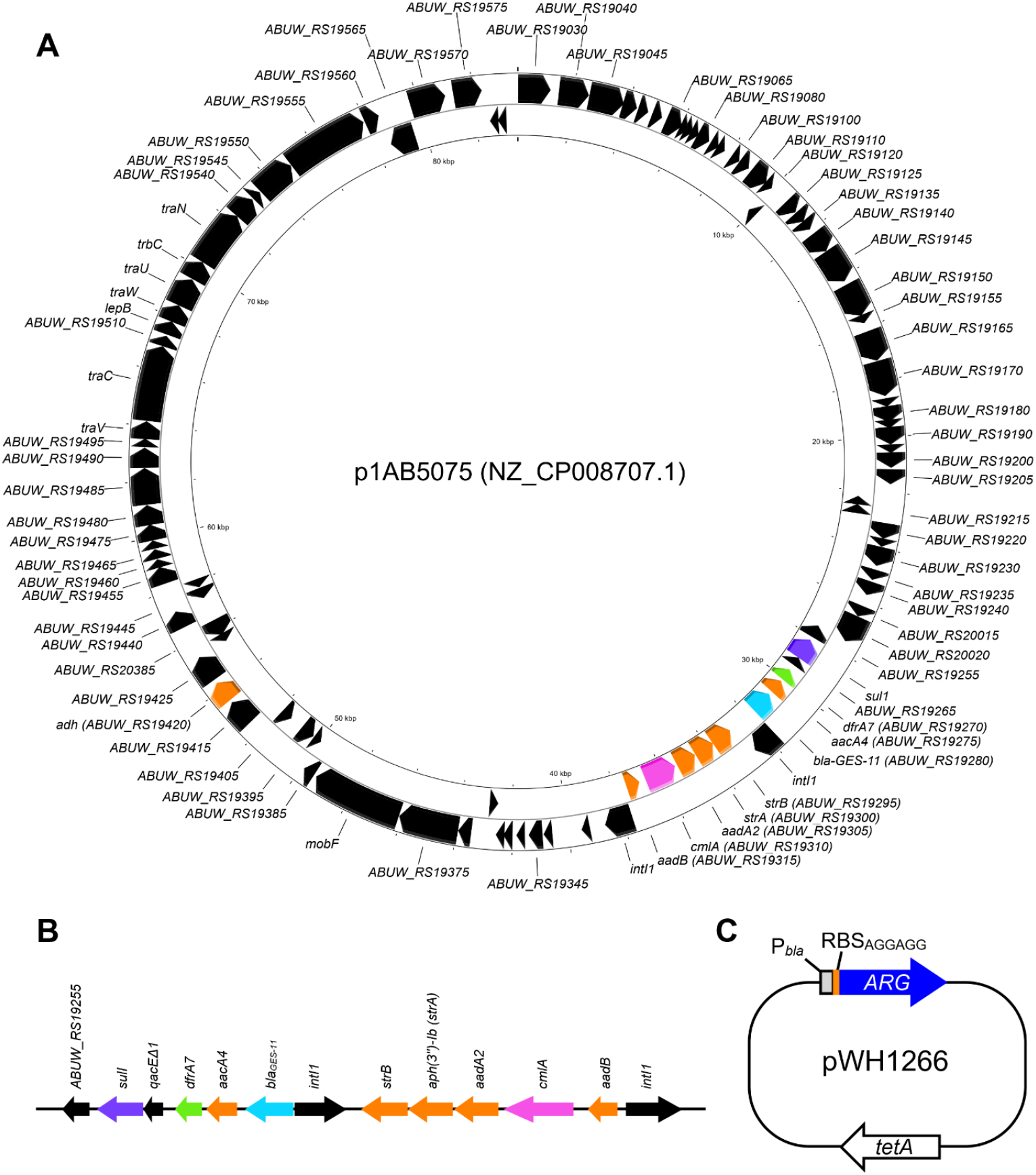
Map of *A. baumannii* AB5075 p1AB5075 (A), Resistance Island-2 (B) and complementation plasmid pWH1266 (C). (A) Aminoglycoside genes are highlighted in orange, chloramphenicol resistance gene *cmlA* in pink, sulphonamide resistance gene *sul1* in light green, beta-lactamase-encoding *bla*_GES-11_ in light blue and trimethoprim resistance gene *dfrA7* in purple. The map was created with Proksee (Grant *et al*., 2023). (B) Linear depiction of p1AB5075-encoded Resistance Island-2. (C) Schematic map of pWH1266 plasmid used for complementation. The *bla* gene of pWH1266 was replaced by the ARG (blue) resulting of transcription of ARGs from the P_*bla*_ promoter (grey, P_*bla*_). All ARGs are translated using the same ribosome binding site (orange, RBS_AGGAGG_). The *tetA* gene of pWH1266 confers resistance to tetracycline. The plasmid is not drawn to scale.

**Figure 2:**
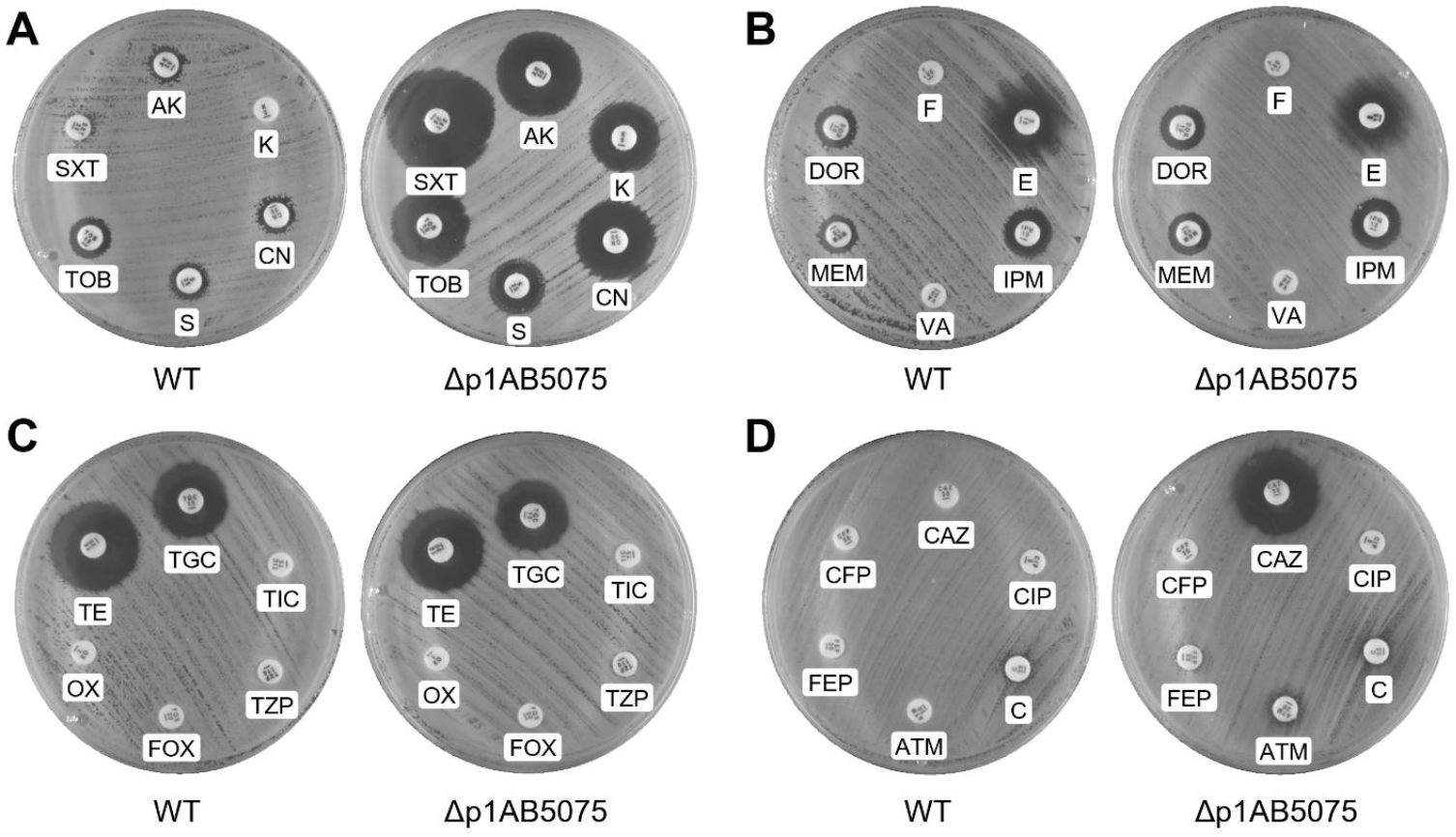
Antibiotic disk diffusion assays comparing *A. baumannii* AB5075 wild-type (WT) and Δp1AB5075 strains. MH2 agar plates were lawned with wild-type *A. baumannii* AB5075 and Δp1AB5075 and antibiotic-containing disks were placed on the agar surface. The plates were incubated for 24 h at 37°C. **(A)** AK (amikacin 30 µg), SXT (trimethoprim-sulfamethoxazole 25 µg), TOB (tobramycin 10 µg), S (streptomycin 25 µg), CN (gentamicin 30 µg), K (kanamycin 5 µg), **(B)** nitrofurantoin (F 100 µg), D (doripenem 10 µg), MEM (meropenem 10 µg), VA (vancomycin 20 µg), IPM (imipenem 10 µg), E (erythromycin 15 µg), **(C)** TGC (tigecycline, 15 µg), TE (tetracycline, 30 µg), OX (oxacillin, 1 µg), FOX (cefoxitin, 30 µg), TZP (piperacillin/tazobactam, 110 µg), TIC (ticarcillin, 75 µg), **(D)** CAZ (ceftazidime, 30 µg), CFP (cefoperazone, 30 µg), FEP (cefepime, 30 µg), ATM (aztreonam, 30 µg), C (chloramphenicol, 30 µg), CIP (ciprofloxacin, 5 µg).

### Plasmid-based complementation of resistance genes reveals individual contributions to AMR

As none of the antibiotic resistance genes have been genetically tested for their individual contributions towards antibiotic resistance in *A. baumannii* AB5075 except for the role of *cmlA* in resistance to chloramphenicol (de Dios *et al*., 2022), we aimed to characterise seven predicted p1AB5075-encoded antibiotic resistance genes *aacA4* (*ABUW_RS19275*), *aadA2* (*ABUW_RS19305*), *aadB* (*ABUW_RS19315*), *adh* (*ABUW_RS19420*), *strA* (*ABUW_RS19300*), *strB* (*ABUW_RS19295*), and *bla*_*GES-11*_ (*ABUW_RS19280*) by ectopically expressing them from a plasmid in the Δp1AB5075 strain. Plasmid-based complementation in *A. baumannii* routinely utilises the shuttle plasmid (or its origin of replication) pWH1266, which has been assembled from pBR322 and pWH1277 (Hunger *et al*., 1990). The latter pWH1277 is only partially sequenced, therefore we first sequenced the plasmid pWH1266, which assembled into a plasmid of 8911 bp. To compare resistance levels, all genes were expressed from the same pWH1266-endogenous promoter of the *bla* gene (**Figure 1C**) and equipped with the same ribosome binding site (AGGAGG) to ensure equal rate of translation. During cloning, the *bla* gene of pWH1266 is removed, leaving *tetA* as the sole (tetracycline) resistance gene. As controls, AB5075 WT and Δp1AB5075 were equipped the pWH1266 “empty” plasmid. Antibiotic disk diffusion assays were performed as before, and the phenotypes in WT and Δp1AB5075 strains remained the same in the presence of pWH1266 compared to the strains without pWH1266 (**Figure 3, Supplementary Figures 1 & 2**). As expected, mild resistance to tetracycline was acquired, which is mediated by *tetA* located on pWH1266 (**Supplementary Figure 2**). Typically, 12.5 µg/ml tetracycline is used in liquid broth, therefore, full resistance is not achieved to the 30 µg of the antibiotic-containing disk. Increased aminoglycoside sensitivity was noted upon deletion of p1AB5075 (Figure 2, Figure 3), and resistance could be restored by expression of either *adh* (**Figure 3C**), *aadB* (**Figure 3D**), *aadA2* (**Figure 3E**), *aacC4* (**Figure 3F**) or *strA* (**Figure 3G**). Expression of *adh* (APH(3’)-VI), which is not part of RI-2 and is flanked by two IS30 family transposases, restored resistance kanamycin and further increased resistance to amikacin in comparison to the wild-type strain (**Figure 3A & C**). Expression of *aadB* (ANT(2’’)-Ia) restored resistance to kanamycin and increased resistance to gentamicin, and tobramycin (**Figure 3 A & D**). Expression of *aacC4* only partially restored resistance to kanamycin (Figure 3F) and *strA* provided resistance to streptomycin (**Figure 3G**). Expression of *strB* did not restore resistance to any of the tested antibiotics (**Figure 3H**). Therefore, the disk diffusion assays detected genes responsible for p1AB5075-mediated aminoglycoside resistance showing also overlap in providing resistance, where *adh, aadB* and to a lower extent *aacCA4* restored resistance to kanamycin (**Figure 3C, D & F**), and *aadA2* and *strA* providing resistance to streptomycin (**Figure 3E & G**). Previously, *aacA4* was described as a potential pseudogene (Gallagher *et al*., 2015), we now show it is functional. The gene outside of RI-2, *adh*, was the only gene providing level resistance to amikacin, while RI-2-encoded *aadB* was the only gene restoring resistance to gentamicin and tobramycin. Expression of *bla*_GES-11_ restored resistance to three antibiotics: ceftazidime, cefepime and aztreonam (**Figure 4**), which was in line with previous work (Moubareck *et al*., 2009).

**Figure 3:**
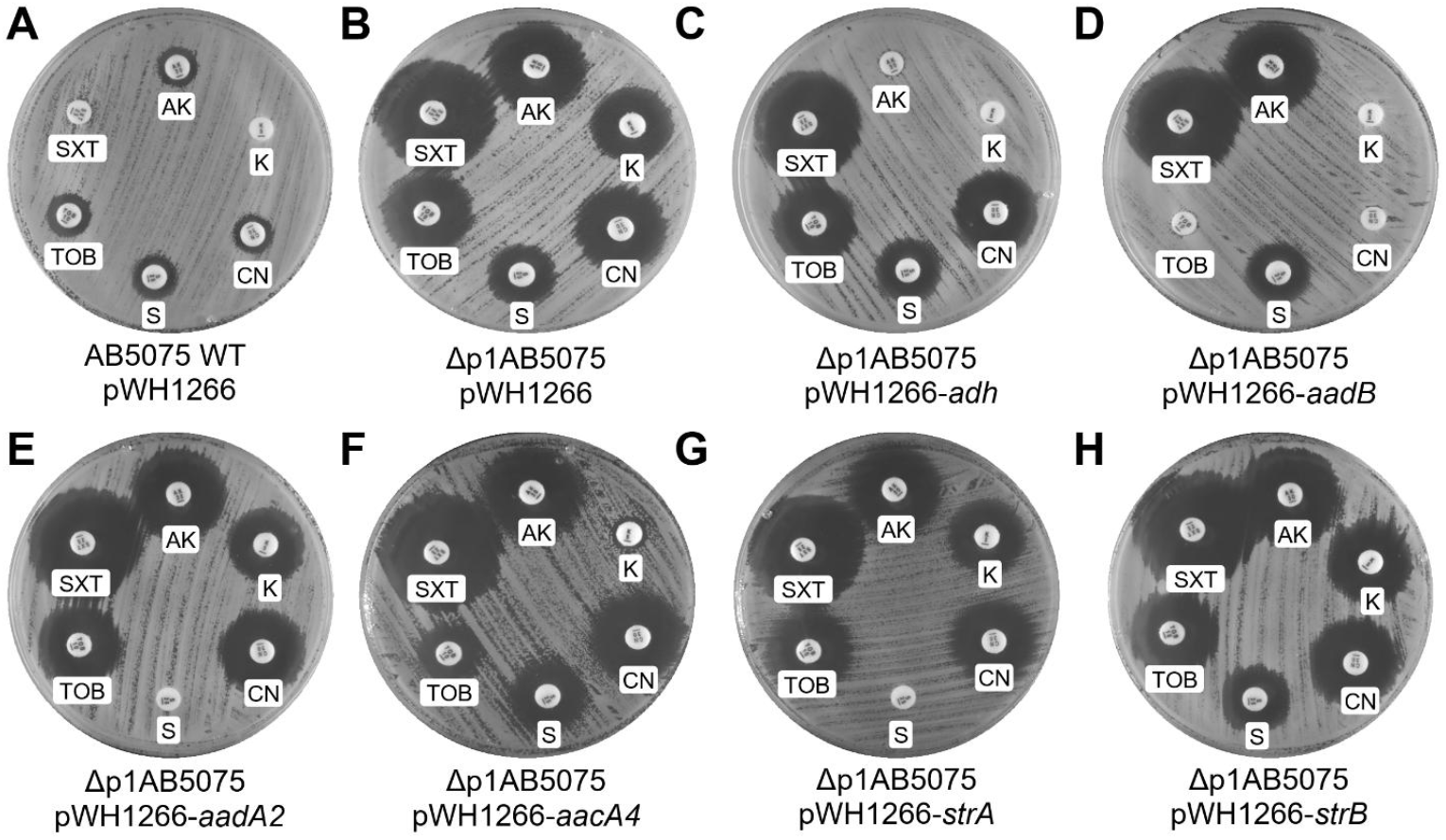
Antibiotic disk diffusion assays comparing *A. baumannii* AB5075 wild-type (WT, A) and Δp1AB5075 (B) strains carrying pWH1266 or pWH1266-ARG: pWH1266-*adh* (C), pWH1266-*aadB* (D), pWH1266-*aadA2* (E), pWH1266-*aacA4* (F), pWH1266-*strA* (G) or pWH1266- *strB* (H). AK (amikacin 30 µg), SXT (trimethoprim-sulfamethoxazole 25 µg), TOB (tobramycin 10 µg), S (streptomycin 25 µg), CN (gentamicin 30 µg), K (kanamycin 5 µg). MH2 agar plates containing tetracycline were lawned with wild-type *A. baumannii* AB5075 and Δp1AB5075 and antibiotic-containing disks were placed on the agar surface. The plates were incubated for 24 h at 37°C.

**Figure 4:**
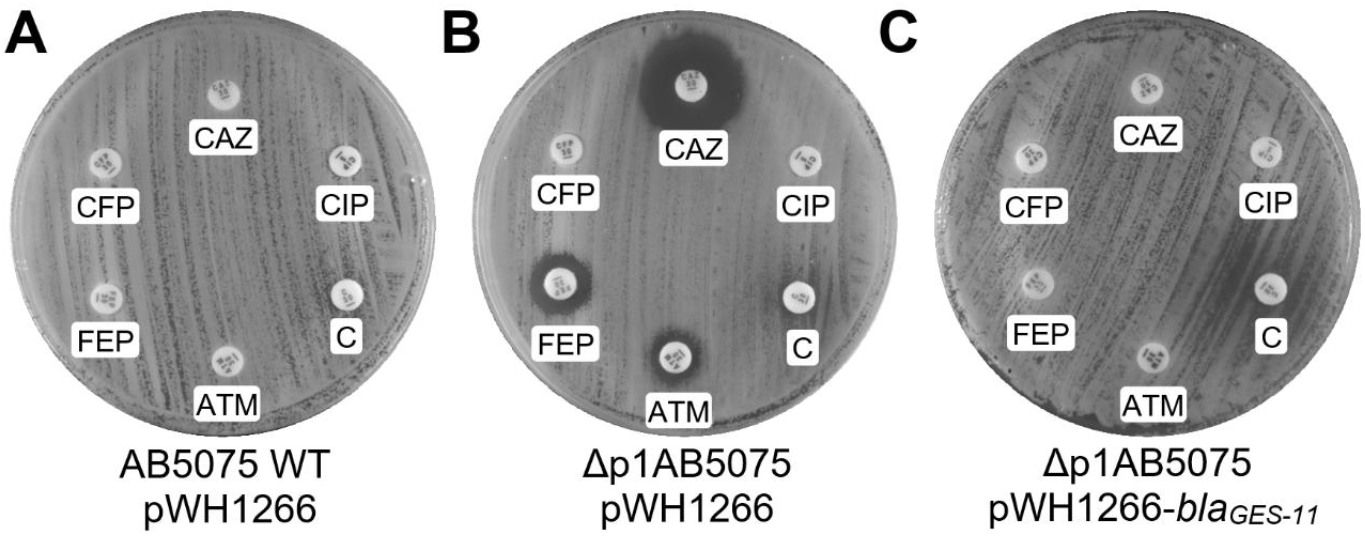
Antibiotic disk diffusion assays comparing *A. baumannii* AB5075 wild-type (WT, A) and Δp1AB5075 (B) strains carrying pWH1266 or pWH1266-*bla*_*GES-11*_ (C). CAZ (ceftazidime, 30 µg), CFP (cefoperazone, 30 µg), FEP (cefepime, 30 µg), ATM (aztreonam, 30 µg), C (chloramphenicol, 30 µg), CIP (ciprofloxacin, 5 µg). MH2 agar plates containing tetracycline were lawned with wild-type *A. baumannii* AB5075 and Δp1AB5075 and antibiotic-containing disks were placed on the agar surface. The plates were incubated for 24 h at 37°C.

To better quantify the contributions of the ARGs to antibiotic resistance and because there was overlap in providing resistance to several antibiotics, minimal inhibitory concentrations were determined by broth microdilution assays for selected antibiotics (**Table 3**). Expression of ARGs from pWH1266 not only restored resistance, but also routinely increased MICs, likely due to overexpression from the *bla* promoter of the pWH1266 plasmid (**Table 3**). Expression of *aadB* increased the MIC to tobramycin from 32 to 128 μg/ml and to gentamicin from 256 to >4096 μg/ml, *adh* increased the MIC to amikacin from 512 to 2048 μg/ml, *aadA2* and *strA* increased the MIC to streptomycin from 1024 to >2048 μg/ml (**Table 3**) matching the data obtained from the disk diffusion assays (**Figure 2, Table 2**). Complementation with pWH1266-*aacA4* restored the resistance to tobramycin to wild-type levels (32 μg/ml) but also provided an increased low-level resistance to multiple aminoglycosides (increases of gentamicin 4-fold, kanamycin 64-fold and amikacin 4-fold) compared to Δp1AB5075 pWH1266 but did not reach wild-type resistance levels. As observed in disk diffusion assays, *strB* did not recover any of the antibiotic sensitivities.

**Table 3:** Minimum inhibitory concentration of aminoglycoside antibiotics determined by broth microdilution assay. Values are in μg/ml. CN=gentamicin, K=kanamycin, AK=amikacin, S=streptomycin, TOB=tobramycin, CEF=cefepime, IMP=imipenem, MEM=meropenem. ND=not determined.

| Strain | CN | K | AK | S | TOB | CEF | IMP | MEM |
| --- | --- | --- | --- | --- | --- | --- | --- | --- |
| WT pWH1266 | 256 | > 4096 | 512 | 1024 | 32 | 256 | 64 | 16 |
| Δp1AB5075 pWH1266 | 8 | 8 | 8 | 256 | 0.5 | 32 | 64 | 8 |
| Δp1AB5075 pWH1266-*aadB* | > 4096 | 2048 | 8 | 256 | 128 | 32 | 64 | 8 |
| Δp1AB5075 pWH1266-*adh* | 8 | >4096 | 2048 | 256 | 0.5 | 32 | 64 | 8 |
| Δp1AB5075 pWH1266-*aadA2* | 8 | 16 | 8 | >2048 | 0.5 | 32 | 64 | 8 |
| Δp1AB5075 pWH1266-*aacA4* | 32 | 512 | 32 | 256 | 32 | 32 | 64 | 8 |
| Δp1AB5075 pWH1266-*strA* | 8 | 16 | 8 | >2048 | 0.5 | 32 | 64 | 8 |
| Δp1AB5075 pWH1266-*strB* | 8 | 16 | 8 | 256 | 0.5 | 32 | 64 | 8 |
| Δp1AB5075 pWH1266-*bla<sub>GES-11</sub>* | ND | ND | ND | ND | ND | 1024 | 64 | 8 |

## DISCUSSION

Multi-drug resistance in *A. baumannii* is mediated through extensive genetic diversity supported by different resistance islands, plasmids, including p1AB5075, and other mobile genetic elements. Here, we characterised the contribution of p1AB5075 and nine p1AB5075-encoded ARGs towards antibiotic resistance. We observed that loss of p1AB5075 renders *A. baumannii* AB5075 sensitive to multiple antibiotics including aminoglycosides, cephalosporins, trimethoprim/sulfamethoxazole and chloramphenicol. We used this Δp1AB5075 genetic background to genetically dissect the contributions of p1AB5075-encoded antibiotic resistance genes. As the plasmid p1AB5075 is transferrable from AB5075 to many other *A. baumannii* and non-*baumannii* strains, transfer would convert any antibiotic sensitive strain to a MDR strain (Nasser *et al*., 2024; Martz *et al*., 2025). We determined that amikacin resistance is predominantly mediated by a gene outside of RI-2 (*adh*, (*ABUW_RS19420*, APH(3’)-VI), which is flanked by IS30 family transposases, which have been described to contain ARGs in *A. baumannii* including the first description of a *bla*_NDM-1_ in Poland (Jachowicz-Matczak *et al*., 2025). However, overexpression of *aacA4* increased the MIC to amikacin four-fold, showing that RI-2 amplification through recombination alone can lead to substantial amikacin resistance (Anderson *et al*., 2020). Indeed, overexpression of *aacA4* conferred low level cross resistance to other aminoglycosides (gentamicin, kanamycin and streptomycin) as well, expanding our knowledge of *A. baumannii* aminoglycoside resistance. Chloramphenicol resistance remained unchanged in the crude disk diffusion assay upon deletion of p1AB5075, which is in agreement with a previous study (de Dios *et al*., 2022), and is due to the presence of addition chloramphenicol resistance genes *craA* and *cpxE* (Karalewitz and Miller, 2018; Foong *et al*., 2019). To our knowledge, although expected upon loss of *bla*_*GES-11*_ the sensitivity to cephalosporins has not been shown for AB5075, yet. We now confirm experimentally that this is due to the presence of *bla*_GES-11_ located in RI-2. AB5075 Δp1AB5075 remains resistant to carbapenems imipenem and meropenem though, due to the presence of an *oxa-23* gene on the chromosome. Similar to AB5075, a clinical isolate of *A. baumannii* isolate from Tunisia with a chromosomally encoded *oxa*-23 and a plasmid-encoded *bla*_GES-11_ copy isolated from a clinical was reported in 2014 (Charfi-Kessis *et al*., 2014). Conjugation of the plasmid carrying *bla*_GES-11_ led to increased resistance to ticarcillin, ticarcillin/clavulanic acid, piperacillin, piperacillin/tazobactam, cefotaxime, ceftazidime, cefepime, aztreonam and even a slight increase to meropenem and imipenem, which might have been masked by the presence of *oxa-23* in our strains (Moubareck *et al*., 2009). A concerning finding was that *bla*_GES-11_ may mutate (Gly170Ser) to *bla*_GES-14_ resulting in a significant increase in imipenem (from 2 to >32 µg/ml) and meropenem (from 4 to 32 µg/ml) resistance as shown in *A. baumannii* isolated from wound infections of soldiers of Ukraine (Kondratiuk *et al*., 2025).

Apart from the clear role of pAB5075 for AMR in AB5075, future studies may investigate the wider impact on *A. baumannii* AB5075 biology as the plasmid is transferrable. Other examples have been reported where (loss) of plasmids had profound effects on gene regulation and pathogenesis. The plasmid pAB3 was shown to repress type-6 secretion in *A. baumannii* ATCC17978 (Weber *et al*., 2015) and a plasmid-encoded copy of H-NS located on plasmid pAB5 in *A. baumannii* UPAB1 was reported to regulate biofilm formation (Benomar *et al*., 2021). As p1AB5075 also contains a copy of an H-NS family protein, there is potential that it might mediate similar crosstalk between p1AB5075- and chromosomally encoded genes. Certainly, a more comprehensive study of plasmids in *A. baumannii* will be required to better understand their impact on *A. baumannii* biology beyond AMR.

## AUTHOR CONTRIBUTIONS

Ideas; formulation or evolution of overarching research goals and aims (OP, KS, CK). Preparation, creation and/or presentation of the published work, specifically visualization/data presentation (OP, CK). Conducting a research and investigation process, specifically performing the experiments, or data/evidence collection (OP, ASE, KS, CK). Oversight and leadership responsibility for the research activity planning and execution, including mentorship external to the core team (KS, CK). Acquisition of the financial support for the project leading to this publication (CK). Preparation, creation and/or presentation of the published work, specifically writing the initial draft (OP, KS, CK). Preparation, creation and/or presentation of the published work by those from the original research group, specifically critical review, commentary or revision – including pre- or post-publication stages (OP, ASE, KS, CK).

## ACKNOWLEDGEMENTS

Deirdre Muldowney (TCD) is acknowledged for technical assistance.

