## Supplementary Material for "Contributions of plasmid p1AB5075-encoded antibiotic resistance genes to multidrug resistance of *Acinetobacter baumannii* AB5075"

**Table S1.** Oligonucleotides used in this study. Red sequences indicate sequences homologous to ends of linearised pWH1266. Purple, underlined sequences indicate ribosome binding site sequence.

| Name | Sequence (5′-3′) | Purpose |
|---|---|---|
| pWH1266_SLiCE_F | TATTGAAGCATTTATCAGGG | Amplification of pWH1266 backbone |
| pWH1266_SLiCE_R | CCTCACTGATTAAGCATTGG | Amplification of pWH1266 backbone |
| *aac(6')-Ib3* F bla-pWH1266 | CCAATGCTTAATCAGTGAGGCAGGGCCATTACCGATTACGCC | Amplification of *aac(6')-Ib3* |
| *aac(6')-Ib3* R bla-pWH1266 | CCCTGATAAATGCTTCAATAAGGAGGGCATCGTGACCAACAGCAACG | Amplification of *aac(6')-Ib3* |
| *aph(3")-lb (StrA)* F bla-pWH1266 | CCAATGCTTAATCAGTGAGGGTCTTCTATAGGTTTCAATCCC | Amplification of *aph(3")-lb* (*strA*) |
| *aph(3")-lb (StrA)* R bla-pWH1266 | CCCTGATAAATGCTTCAATAAGGAGGCTCCATTGAATCGGACTTATAT | Amplification of *aph(3")-lb* (*strA*) |
| *aph(6)-l (StrB)* F bla-pWH1266 | CCAATGCTTAATCAGTGAGGCCGCTGCTATAGGGGTC | Amplification of *aph(6)-l* (*strB*) |
| *aph(6)-l (StrB)* R bla-pWH1266 | CCCTGATAAATGCTTCAATAAGGAGGGGTTGATGTTCATGCCGC | Amplification of *aph(6)-l* (*strB*) |
| *aadA1* F bla-pWH1266 | CCAATGCTTAATCAGTGAGGCCACGTCGAAAAACAAAATCAC | Amplification of *aadA1* |
| *aadA1* R bla-pWH1266 | CCCTGATAAATGCTTCAATAAGGAGGGACATCATGAGGGTAGCGG | Amplification of *aadA1* |
| *aadB* F bla-pWH1266 | CCAATGCTTAATCAGTGAGGCTGCTGGCTATCTCATGATTG | Amplification of *aadB* |
| *aadB* R bla-pWH1266 | CCCTGATAAATGCTTCAATAAGGAGGGCCGCATGGACACAAC | Amplification of *aadB* |
| *aph(3')-VI* F bla-pWH1266 | CCAATGCTTAATCAGTGAGGCTCAAGCATTAATGCAGTACGATC | Amplification of *aph(3')-VI* |
| *aph(3')-VI* R *bla*-pWH1266 | CCCTGATAAATGCTTCAATAAGGAGGACTTGATGGAATTGCCCAATATTA | Amplification of *aph(3')-VI* |
| *bla$_{GES}$ 11* F *bla*-pWH1266 | CCAATGCTTAATCAGTGAGGGCCTGAGTTAAGCCGCGGTGC | Amplification of *bla$_{GES}$ 11* |
| *bla$_{GES}$ 11* R *bla*-pWH1266 | CCCTGATAAATGCTTCAATAAGGAGGTCACCATGCGCTTCATTCACGCAC | Amplification of *bla$_{GES}$ 11* |

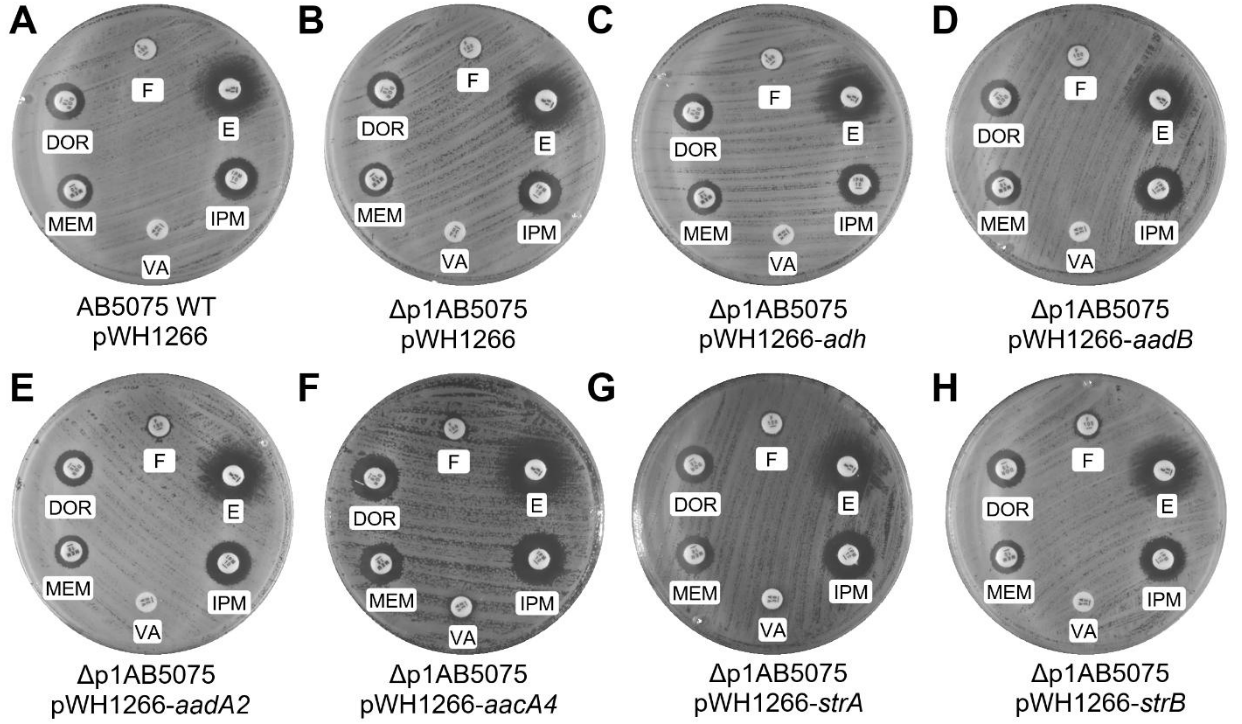


**Supplementary Figure 1: Antibiotic disk diffusion assays comparing *A. baumannii* AB5075 wild-type (WT, A) and Δp1AB5075 (B) strains carrying pWH1266 or pWH1266-ARG:** pWH1266-*adh* (C), pWH1266-*aadB* (D), pWH1266-*aadA2* (E), pWH1266-*aacA4* (F), pWH1266-*strA* (G) or pWH1266-*strB* (H). Nitrofurantoin (F, 100 µg), erythromycin (E, 15 µg), imipenem (IMP 10 µg), vancomycin (VA, 30 µg), meropenem (MEM, 10 µg), doripenem (DOR, 10 µg). MH2 agar plates containing tetracycline were lawned with wild-type *A. baumannii* AB5075 and Δp1AB5075 and antibiotic-containing disks were placed on the agar surface. The plates were incubated for 24 h at 37°C.

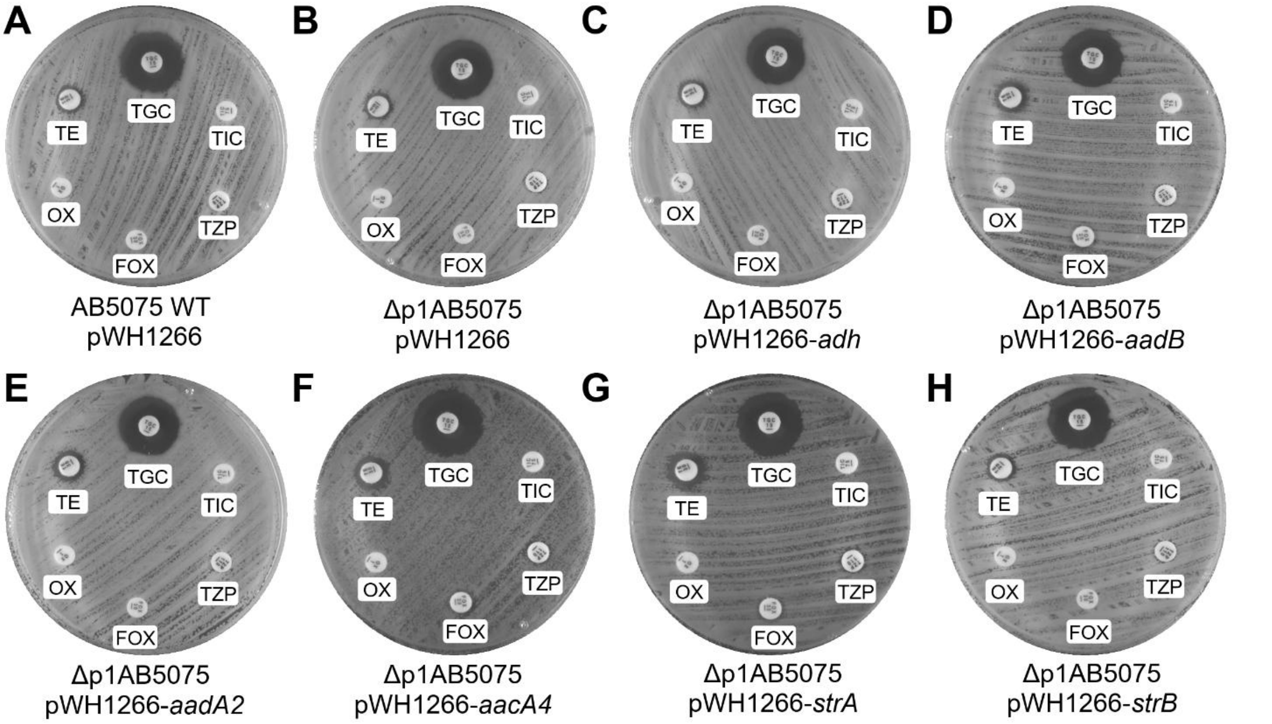


**Supplementary Figure 2: Antibiotic disk diffusion assays comparing *A. baumannii* AB5075 wild-type (WT, A) and Δp1AB5075 (B) strains carrying pWH1266 or pWH1266-ARG:** pWH1266-*adh* (C), pWH1266-*aadB* (D), pWH1266-*aadA2* (E), pWH1266-*aacA4* (F), pWH1266-*strA* (G) or pWH1266-*strB* (H). Tigecycline (TGC 15 µg), ticarcillin (TIC, 75 µg), piperacillin/tazobactam (TZP, 110 µg), cefoxitin (FOX, 30 µg), oxacillin (OX, 1 µg), tetracycline (TE, 30 µg). MH2 agar plates containing tetracycline were lawned with wild-type *A. baumannii* AB5075 and Δp1AB5075 and antibiotic-containing disks were placed on the agar surface. The plates were incubated for 24 h at 37°C.
